# Lactylation landscape of mitochondrial proteins in myocardial infarction

**DOI:** 10.64898/2026.04.27.718938

**Authors:** Ashlesha Kadam, Shiridhar Kashyap, Kunal Samantaray, Natasha Jaiswal, Shanikumar Goyani, Philip A Kramer, Pourhadi Hadi, Jingyun Lee, Cristina M Furdui, Pooja Jadiya, Dhanendra Tomar

**Author notes:** These authors share corresponding authorship. **Correspondence**: **Dhanendra Tomar, PhD, FAHA,** Assistant Professor, Department of Cardiovascular Medicine, Wake Forest University School of Medicine, Medical Center Blvd. NRC/Commons Building: Office 219, Winston-Salem, NC 27157 USA, | https://tomarlab.org, **Pooja Jadiya, PhD,** Assistant Professor, Section of Gerontology and Geriatric Medicine, Department of Internal Medicine, Wake Forest University School of Medicine, Medical Center Blvd. NRC/Commons Building: Office 220, Winston-Salem, NC 27157 USA, | https://jadiyalab.org.

## Abstract

Metabolic reprogramming is a hallmark of myocardial infarction (MI), in which cardiomyocytes shift from fatty acid oxidation to anaerobic glycolysis, leading to elevated lactate production and mitochondrial dysfunction. Lactylation, a recently described lysine post-translational modification, has emerged as a metabolic signaling mechanism; however, its role within mitochondria during MI remains poorly understood. Here, we define the mitochondrial lactylome following MI and examine how modulation of lactate transport influences mitochondrial metabolism and redox homeostasis. Using quantitative proteomics, we identify extensive remodeling of mitochondrial protein lactylation after MI, affecting enzymes involved in bioenergetics, redox regulation, and metabolic control. Pharmacological inhibition of monocarboxylate transporter-1 (MCT1) using AZD3965 further reshapes the mitochondrial lactylome, increasing lactylation of specific metabolic and redox-associated proteins without uniformly exacerbating mitochondrial dysfunction. Despite sustained impairment of global cardiac function, MCT1 inhibition attenuates post-MI fibrosis and inflammation and partially restores mitochondrial respiratory capacity. Consistent with in vivo findings, genetic or pharmacological inhibition of MCT1 in hypoxic cardiomyocytes-derived cells reduces mitochondrial reactive oxygen species, decreases inhibitory pyruvate dehydrogenase phosphorylation, and improves mitochondrial bioenergetics. Together, these findings reveal that mitochondrial lactylation is a context-dependent regulator of mitochondrial metabolism and redox balance following MI. Rather than acting solely as a pathological modification, lactylation integrates lactate availability with mitochondrial function to influence inflammatory and fibrotic remodeling, highlighting mitochondrial metabolic plasticity as a potential therapeutic target in ischemic heart disease.

**Highlights:**

- Myocardial infarction (MI) increases mitochondrial protein lactylation, with 361 identified lactylated proteins.
- AZD3965-mediated MCT1 inhibition further elevates mitochondrial lactylation.
- Distinct alterations in mitochondrial proteins and pathways (TCA cycle, amino acid metabolism, gene expression) were observed.
- AZD3965 reduces cardiac fibrosis and inflammation and partly improves mitochondrial respiration post-MI, but cardiac function remains impaired.

Graphical Abstract

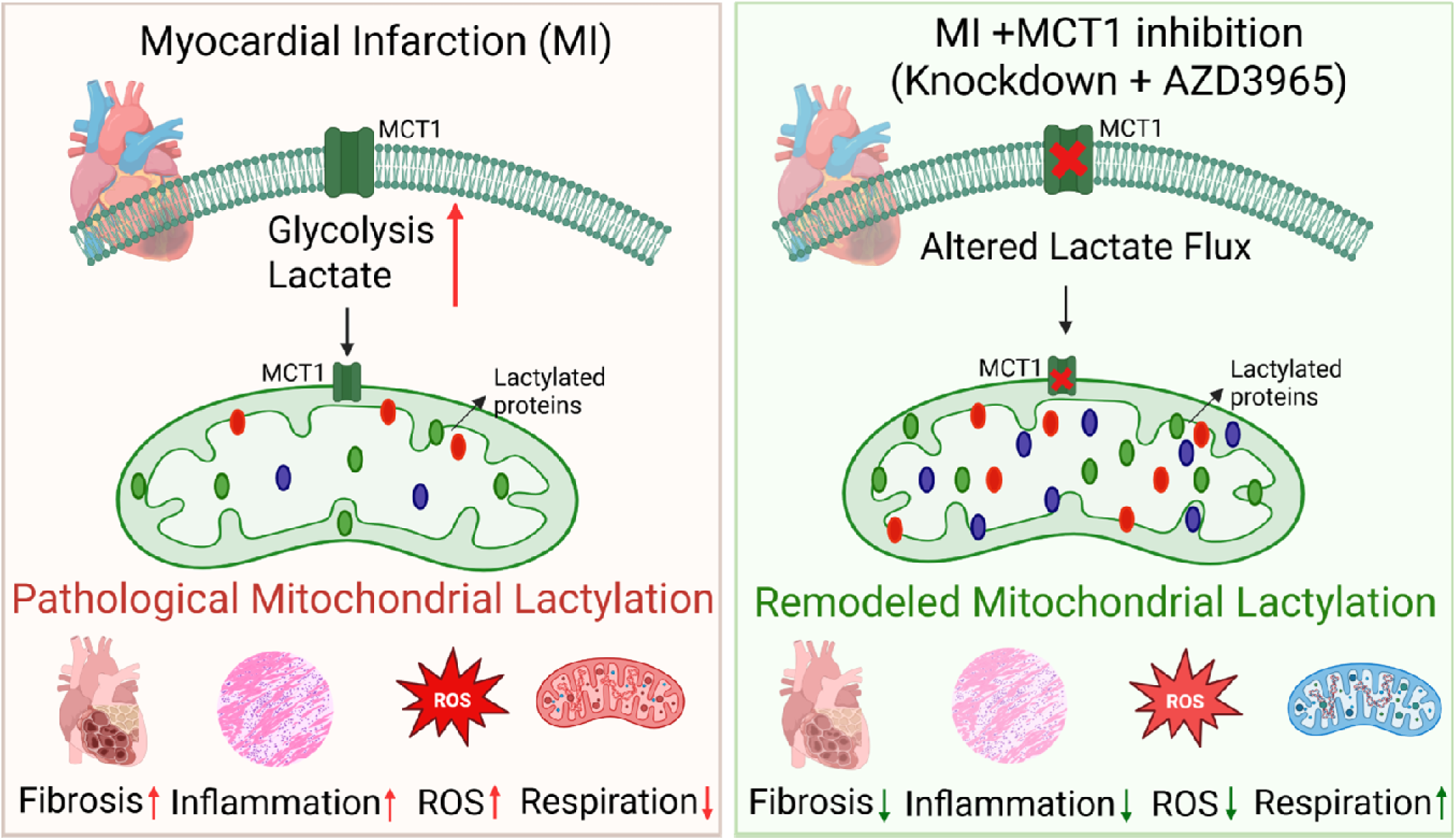

## Introduction

Myocardial infarction (MI) is a major contributor to illness and death, arising from the sudden blockage of oxygen and nutrients to cardiac tissue caused by coronary artery occlusion [1]. This ischemic event leads to cellular dysfunction, including oxidative stress, depletion of ATP, ion imbalances, and ultimately death of cardiomyocytes. The resulting necrotic tissue is replaced by fibrotic scarring, which drives long-term cardiac remodeling and impaired function [1]. Mitochondria are central to the fate of cardiomyocytes, shifting from energy production to initiating cell death pathways under stress [2]. Essential mitochondrial processes, including bioenergetics, dynamics, and mitophagy, are tightly regulated by post-translational modifications (PTMs) to maintain cellular homeostasis. A defining feature of MI is metabolic reprogramming, where cardiomyocytes transition from fatty acid oxidation to anaerobic glycolysis, resulting in an accumulation of lactate [3]. Lactate is transported across membranes by monocarboxylate transporters (MCT1–4), which help regulate intracellular pH and lactate balance [4].

Lactylation, a recently discovered post-translational modification targeting lysine residues, was initially identified on histones [5]. It is now recognised on non-histone proteins, suggesting broader roles in gene regulation and cellular adaptation to metabolic stress [6]. Given that mitochondria are both major regulators of lactate production and primary determinants of cellular bioenergetic capacity, mitochondrial proteins represent particularly compelling targets for lactylation. Lactate has been proposed to be directly oxidized by mitochondria, a model known as the intracellular lactate shuttle (ILS) [7]. Energized mitochondria can produce and export lactate, using it as a rapid means to regulate matrix redox state and ROS, thereby positioning lactate as a mitochondrial stress-buffering output rather than merely a metabolic fuel [8]. Based on the existing literature, mitochondrial proteins account for a substantial fraction of lactylation targets across species, including 5.5% in HEK293 [9], 6.7% in mouse brain [10], 11% in *T. brucei [11]*, 27% in *B. Cinerea [12]*, and 9.1% in rice [13], collectively highlighting mitochondria as a major hub for protein lactylation. The role of lactylation in disease progression associated with mitochondrial dysfunction is now emerging [6], its specific role in MI remains largely unclear. Elevated lactate levels, an outcome of metabolic reprogramming, are linked to increased mortality in heart failure [14]. These levels promote cardiac fibrosis by stimulating the endothelial-to-mesenchymal transition, mediated through lactylation of transcription factors like Snail1 [15], highlighting the pathological significance of this modification. Nevertheless, the extent of mitochondrial protein lactylation following MI and its relationship with mitochondrial activity, particularly in the context of energy metabolism and redox regulation, remains unexplored. This knowledge gap is critical, given that mitochondria play a central role in regulating cellular metabolism, reactive oxygen species (ROS) production, and cardiomyocyte survival during ischemic stress. Targeting lactylation-dependent mitochondrial pathways may therefore offer a promising approach to mitigate mitochondrial dysfunction and adverse cardiac remodeling following MI. Our study aims to address this gap by profiling mitochondrial lactylation after MI and evaluating the impact of inhibiting Monocarboxylate Transporter 1 (MCT1) which fluxes lactate to mitochondrial matrix. We investigate the effect of AZD3965, a selective MCT1 inhibitor that increases cytosolic lactate by blocking its bidirectional transport [4]. Recent research shows that blocking MCT1 with AR-C155858 (a dual MCT1/MCT2 inhibitor) in perfused mouse hearts led to retention of succinate in cells, exacerbating ROS generation and ischaemia-reperfusion (IR) injury [16]. AZD3965 similarly inhibits flux of lactate through the MCT1, resulting in increased intracellular lactate and triggering context-dependent metabolic changes in the mitochondria of cells expressing MCT1 [4, 17, 18]. This can either inhibit further glycolysis through feedback mechanisms, forcing cells to depend on alternative metabolic pathways, or increase the conversion of intracellular lactate back to pyruvate, which is then redirected towards mitochondrial oxidation as a compensatory mechanism. AZD3965, with broader specificity and greater potency for MCT1, is currently undergoing clinical trials due to its improved properties compared to its predecessor, AR-C155858 [19–21]. Therefore, exploring the effects of AZD3965 as a mediator for mitochondrial lactylation may provide important mechanistic insight into lactate-driven metabolic and redox remodeling during MI. Our findings provide the first comprehensive profiling of mitochondrial lactylation following MI and lay the groundwork for future mechanistic and therapeutic investigations.

## Methods

### Animals

All animal experiments were conducted in accordance with the Wake Forest University School of Medicine Institutional Animal Care and Use Committee (IACUC) guidelines. Adult male C57BL/6NJ mice (10–12 weeks old) underwent MI induction by permanent ligation of the left anterior descending (LAD) coronary artery, with successful infarction confirmed by echocardiography. Mice were randomly assigned to experimental groups and housed under standard laboratory conditions. AZD3965 (MedChemExpress, Cat# HY-12750), a selective MCT1 inhibitor, was administered by oral gavage at 25 mg/kg beginning 24 h prior to surgery and continued daily for two weeks (Fig. S1A). Sham and MI control groups received vehicle alone (10% DMSO, 90% corn oil). At the end of the experimental period, animals were euthanized, and blood and heart tissues were collected for downstream analyses.

### Cell culture, transfection, and chemical treatments

AC16, an immortalized human cardiomyocyte-derived cell lines, were cultured in DMEM/F12 medium (Corning, Cat# 10-090-CV) supplemented with 10% fetal bovine serum (Gibco, Cat# 26140079) and 1% penicillin/streptomycin (Corning, Cat# 30-002-0CI) at 37 °C in 5% CO₂. For pharmacological inhibition of MCT1, cells were treated with AZD3965 (25 nM) for 3 h. For genetic knockdown, cells were transfected with control or MCT1-specific siRNA (Origene, Cat# SR321815) using Lipofectamine RNAiMAX (Invitrogen, Cat# 13778-150), and experiments were performed 36 h post-transfection. For imaging studies, cells were plated on poly-D-lysine-coated glass-bottom dishes (MatTek, Cat# P35GC). Hypoxia was induced by incubating cells in a controlled hypoxia chamber (1% O₂, 5% CO₂, 94% N_2_) in serum-free DMEM for 4 h[22, 23]. For plate-reader-based assays, chemical hypoxia was induced using CoCl₂ (800 µM, 4 h) [24].

### Histological analysis

Hearts were fixed in 4% paraformaldehyde overnight, paraffin-embedded, and sectioned at 4 µm thickness. Sections were stained with Masson’s trichrome (Sigma-Aldrich, Cat#HT15), Sirius Red (Polyscience, Cat#24901), or hematoxylin and eosin (H&E) staining (Fischer Scientific, Cat#NC1470670) following standard protocols [25]. Images were acquired using a Keyence BZ-X700 microscope and quantified using Fiji ImageJ. Multiple sections per heart were analyzed, with representative fields selected from infarct, border, and remote zones. Fibrosis was quantified in infarct, border, and remote zones as percentage area of collagen-positive staining. Inflammation scores were calculated as the ratio of immune cells to total cardiomyocytes [25].

### Echocardiography

Cardiac function was assessed at baseline and at one– and two-weeks post-MI. Mice were anesthetized with isoflurane (induction 2–3%, maintenance 1.8–2%) and imaged using B-mode long-axis and M-mode short-axis views. The mice were placed in a supine position, and body temperature and heart rate (typically 400-500 bpm) were maintained throughout the procedure. B-mode long-axis and M-mode short-axis images of LV were captured. The M-mode was directed through the centre of the 2D parasternal short axis, just distal to the mitral valve leaflets, and the M-mode gains were adjusted to optimize endocardial and epicardial interfaces. Left ventricular dimensions, ejection fraction, fractional shortening, and cardiac output were calculated using standard methods.

### High-resolution respirometry

Mitochondrial respiration was measured in saponin-permeabilized (Thermo Fisher Scientific, Cat#A18820-14; 50 µg/mL for 30 min) heart tissue using an Oroboros O2k Oxygraph [26, 27]. Experiments were performed in the Core for Cellular Respirometry (CCR) shared resource at Wake Forest University School of Medicine. Substrate-uncoupler-inhibitor titration protocols were performed using pyruvate (2 M)/malate (0.8 M) (Sigma-Aldrich, Cat#P2256-5G; MP biomedicals, Cat#521807310), ADP (10 mM & 0.5 M) (MilliporeSigma, Cat#A2754), glutamate (2 M)/succinate (1 M) (Fisher Bioreagents, Cat#BP336-500; MilliporeSigma, Cat#1.06445), FCCP (1 mM) (Sigma-Aldrich, Cat#C2920) and Antimycin A (5 mM) (MilliporeSigma, Cat#A8674). Respiration rates were normalized to tissue wet weight. Cytochrome c (4 mM) (Sigma-Aldrich, Cat#C7752) addition served as a quality control step, with >15% increase indicating compromised mitochondrial integrity.

### Lactylome profiling by LC-MS/MS

Mitochondria were isolated from heart tissues using mitochondria isolation buffer (MIB) to ensure sample quality and purity. The mitochondrial pellet was lysed in RIPA buffer (MilliporeSigma, Cat#20-188) containing Protease Inhibitor Cocktail (Roche, Cat#04693132001). The lysate was clarified by centrifugation at 16,500 g for 20 min. The resulting supernatants were incubated overnight with anti-L-Lactyllysine antibody (PTM Bio, Cat#PTM-1401RM). The immune complexes formed were subsequently incubated for 4 h with protein A-conjugated magnetic beads (Thermo Fisher Scientific, Cat#10002D) at 4°C to pull down bead-bound lactylated proteins. Beads with bound proteins were washed with RIPA buffer containing 2% SDS and used for proteomic analysis to identify and quantify lactylated proteins. The beads were washed three times with 50 mM ammonium bicarbonate, followed by overnight incubation with a digestion solution containing sequencing-grade modified trypsin in 50 mM ammonium bicarbonate. The peptide mixture was purified using a C18 tip, and samples were analysed on an LC-MS/MS system consisting of an Orbitrap Eclipse Mass Spectrometer and a Vanquish Neo nano-UPLC system (Thermo Scientific, Waltham, MA). The peptide mixture was injected via a trap-and-elute procedure using a Thermo Scientific Acclaim PepMap 100 trap column (Cat#164946) and a Thermo Scientific DNV PepMap Neo (Cat#DNV75500PN) analytical column. Peptides were separated using linear gradient elution, consisting of water (A) and 80% acetonitrile (B), both containing 0.1% formic acid. Data were collected by data-independent acquisition (DIA), wherein fragmentation occurred in 44 static DIA windows, each at a width of 10 Th, spanning 400–900 m/z. Cycle time was set to 3 sec between adjacent survey spectra. For protein identification, spectra were searched against the UniProt mouse protein FASTA database (17,082 annotated entries, Oct 2021) using the Pulsar search engine within Spectronaut v20 (Biognosys AG, Switzerland). Search parameters included: FT-trap instrument; parent mass error tolerance, 10 ppm; fragment mass error tolerance, 0.6 Da (monoisotopic); enzyme, trypsin (full); maximum missed cleavages, 2; and variable modifications, +15.995 Da (oxidation) on methionine.

### Lactylome profiling: Data processing and differential abundance analysis

Protein-level abundance matrices were exported and processed in R. Due to the sparsity inherent to affinity-based proteomics, missing values were imputed using a column-wise minimum approach. For each sample, missing entries were replaced with one-fifth of the minimum observed intensity within that sample column, approximating values below the detection limit. Imputation was performed across all quantitative sample columns following numeric coercion and imputed values were used exclusively for downstream analyses. A summary of imputation parameters, including per-column minima, replacement values, and number of imputed entries, was recorded. The processed dataset and imputation metadata were exported as multi-sheet Excel files for downstream analysis. Differential protein abundance was assessed using the linear modeling framework implemented in the limma package. Log2-transformed protein intensities (with a small offset of 1 × 10⁻□) were modeled using a design matrix encoding experimental groups: Sham, MI and MI treated with the MCT1 inhibitor AZD3965.

All pairwise ordered contrasts between groups were evaluated using empirical Bayes moderation (trend and robust enabled) to stabilize variance estimates across proteins. Proteins were included in comparisons only if at least two valid observations were present in each group. Significantly altered proteins were defined by a threshold of p < 0.05 and |log2 fold change| > 1. Benjamini–Hochberg adjusted p-values were calculated and reported but not used for primary filtering to retain sensitivity in pulldown-based datasets. To focus on mitochondrial proteins, differential expression outputs were filtered using a curated mouse mitochondrial gene set (MitoCarta) [28]. In MATLAB, gene identifiers were standardized to uppercase and matched against the reference list. Duplicate entries were removed, and only genes present in both datasets were retained. Filtered mitochondrial subsets and unmatched genes were exported as separate tables for each comparison. Functional enrichment was performed in R using clusterProfiler and ReactomePA. Significant genes were split into upregulated and downregulated sets (p<0.05, |log2FC| ≥ 1), mapped to ENTREZ identifiers, and analyzed for Gene Ontology Biological Processes (GO:BP). Results were filtered at q<0.05 and exported as multi-sheet Excel files. Dot plots of top enriched GO terms were generated for visualization. For each contrast, results were exported as Excel files containing protein annotations, sample-level abundances, log2 fold changes, raw p-values, and adjusted p-values. All analyses were performed using reproducible R and MATLAB scripts incorporating automated sample detection, contrast generation, and batch export of results. R programming and SRplot were used for the chord plot and GO enrichment analysis [29], respectively. Heat map visualization was performed using Morpheus (https://software.broadinstitute.org/morpheus), developed by the Broad Institute.

### Mitochondrial bioenergetics

Cells were subjected to Oxygen Consumption Rate (OCR) measurement at 37□°C in an XF96 extracellular flux analyzer (Seahorse Bioscience). AC16 cells (2□×□10^4^) were plated in XF media pH 7.4 supplemented with 2 mM Glutamine, 25□mM glucose, and 1□mM sodium pyruvate and sequentially exposed to oligomycin (1.5□µM), FCCP (2□µM), and rotenone plus antimycin A (2□µM). Quantification of basal respiration (base OCR – non-mito respiration (post-Rot/AA), ATP-linked respiration (post-oligo OCR−base OCR), and Max respiratory capacity (post-FCCP OCR−post-Rot/AA) was performed.

### Quantitative Real-Time PCR (qRT-PCR) analysis

Total RNA was extracted using the RNeasy Kit (Qiagen, Cat# 74106). Briefly,1 μg of RNA was used to synthesize cDNA using the SuperScript™ IV First-Strand Synthesis System with ezDNase™ Enzyme (Thermo Fisher Scientific, Cat#18091150). qRT-PCR analysis followed manufacturer instructions using PowerUp^TM^ SYBR^TM^ Green master mix (Thermo Fisher Scientific, Cat#A25742). *GAPDH* and *ACTIN* were used as an internal control to normalize RNA. Each sample was run in triplicate, and relative gene expression was analyzed using the 2^^ΔΔCt^ method. A list of all primer sequences is available in the supplementary table S1.

### Evaluation of mitochondrial ROS generation

To measure mitochondrial ROS, AC16 cells (2□×□10^5^) were seeded in MatTek collagen-coated glass-bottom 35 mm culture plates and loaded with 5□μM MitoBright ROS Deep Red (MBR Deep Red) (Dojindo Molecular Technologies, Cat#MT16-12) for 30□min at 37□°C, and measured the fluorescence at 540/ex and 670/em using an Olympus Evident FV4000 Confocal Microscope (Olympus) using a 60x oil objective. The images were quantified as mean fluorescence intensity per cell using ImageJ software, ensuring consistent thresholding and background subtraction across all samples.

### Evaluation of mitochondrial membrane potential

To assess mitochondrial membrane potential, 2×10^5^ AC16 cells were seeded in MatTek collagen-coated glass-bottom 35 mm culture plates and stained with 20 nM TMRM red (Biotium, Cat#70017) for 30 min at 37° C. The cells were imaged using an Olympus Evident FV4000 Confocal Microscope (Olympus) using a 60x oil objective. The mean fluorescence intensity per cell was calculated using ImageJ by subtracting the background intensity.

### Immunoprecipitation and western blotting

Cells were lysed in 1× RIPA lysis buffer supplemented with 1 x protease inhibitor cocktail (Thermo Fischer Scientific, Cat#78429). Cell-lysates were centrifuged, and protein concentration was calculated using the Pierce 660 nm Protein Assay. An equal amount of protein was resolved by SDS–PAGE, transferred to PVDF membranes, blocked, and incubated with primary antibodies against MCT1 (Santa Cruz Biotechnology, Cat#sc-365501), p-PDH (Proteintech, Cat#29580-1-AP), ATP5A1 (Proteintech, Cat#66037-1-Ig), Cytochrome-c (Santa Cruz Biotechnology, Cat#sc-13156 AF680), HSP60 (Proteintech, Cat#66041-1-Ig), TOMM20 (Proteintech, Cat#11802-1-AP), GAPDH (Cell Signalling Technology, Cat#2118S) and pan-lactyllysine (PTM Biolabs, Cat#PTM1401) followed by IRDye-conjugated secondary antibodies. Band intensities were detected using a Bio-Rad ChemiDoc system, quantified with ImageJ, and normalized to ATP5A1 and GAPDH. For endogenous immunoprecipitation, equal amounts of cell lysate protein were incubated overnight with anti-PDK1 (Proteintech, Cat#18262-1-AP) and anti-PDK2 (Proteintech, Cat#15647-1-AP) antibodies (4 µg), followed by capture with Protein A Dynabeads (Thermo Fisher Scientific, Cat#10002D) for 4 h. Rabbit IgG isotype (Thermo Fisher Scientific, Cat#02-6102) was used as a negative control. Input samples were collected before antibody incubation. Immunocomplexes were washed with RIPA buffer containing 2% SDS, eluted in Laemmli buffer, resolved by SDS–PAGE, and then analyzed by western blotting. Protein band intensities were analyzed using ImageJ and normalized to loading controls.

### Submitochondrial protein localization assay

Mitochondria were isolated as described earlier[30]. Briefly, cells were washed with PBS and resuspended in isotonic mitochondria isolation buffer (MIB) [10 mM HEPES (pH 7.5), containing 200 mM mannitol, 70 mM sucrose, and 1 mM EGTA]. Cell suspensions were homogenized by Dounce homogenizer and centrifuged at 500g for 10 min at 4°C. Supernatants were collected and centrifuged at 12,000g for 15 min at 4°C to obtain crude mitochondrial pellets. Pellets were resuspended in mitochondria isolation buffer and washed twice using the spin at 12,000g for 15 min at 4°C. Mitochondrial pellets were resuspended in MIB and digested with proteinase K (3 μg/ml) for 10 min on ice. Proteinase K digestion was stopped by adding the 2 mM PMSF, 1x Protease Inhibitor Cocktail, and 2× SDS loading dye. The protein samples were then heated at 95°C for 10 min and analyzed by western blotting for lactylation.

### Statistical analyses

Statistical analysis was performed using GraphPad Prism 10 (GraphPad Software). Results are represented as mean±standard error. P-value analyses were performed using an unpaired, 2-tailed t-test (for 2 groups) with Welch’s correction. For grouped analyses, 2-way ANOVA with Tukey post-hoc analysis or Dunnett’s multiple comparisons test was applied. P values less than 0.05 (95% confidence interval) were considered significant.

## Results

### MI alters the mitochondrial lactylome

To systematically profile mitochondrial protein lactylation in mouse hearts following MI, heart tissues were harvested two weeks post-MI and subjected to mitochondrial isolation. Purified mitochondrial fractions were processed for denaturing immunoprecipitation, and lactylated proteins were enriched using anti-L-Lactyllysine antibodies prior to mass spectrometry analysis (Fig. 1A, Fig. S1A). Pos-MI survival rates were monitored across experimental groups, revealing no significant differences during the study period (Fig. S1B). Global lactylome profiling identified a total of 1,368 lactylated proteins, of which 361 were annotated as mitochondrial proteins based on MitoCarta classification. Comparative analysis across sham, MI, and MI + AZD3965 groups revealed significant alterations in mitochondrial protein lactylation, defined by an absolute fold change ≥ 2 and P < 0.05 (Fig. 1B, C). Lactylation site prediction using DeepKla software [31]. Further annotated candidate lysine lactylation sites in proteins altered in MI compared with sham controls (Table S2). Volcano plot analysis demonstrated distinct patterns of mitochondrial protein lactylation following MI (Fig. 1B). Specifically, lactylation was significantly reduced in 21 proteins, including PUS1, NDUFS7, SHMT2, FKBP10, HSPE1, NDUFB10, MAOA, PCK2, HINT1, C1QBP, MTHFD2, SOD1, SND1, DCAKD, TOMM22, PDK2, CYB5B, CHCHD2, ACOT9, FASN, and GFM1. In contrast, lactylation was increased in 18 proteins, including ACADM, SLC25A4, SLC25A3, SLC25A5, SLC25A12, PCCA, SQOR, VDAC2/3, UQCRC1/2, HADHA, GATD3A, TUFM, CPT1B, BDH1, POLB, and CKMT1. Heatmap visualization further highlighted distinct and reproducible changes in mitochondrial protein lactylation between sham and MI groups (Fig. 1C).

**Figure 1.**
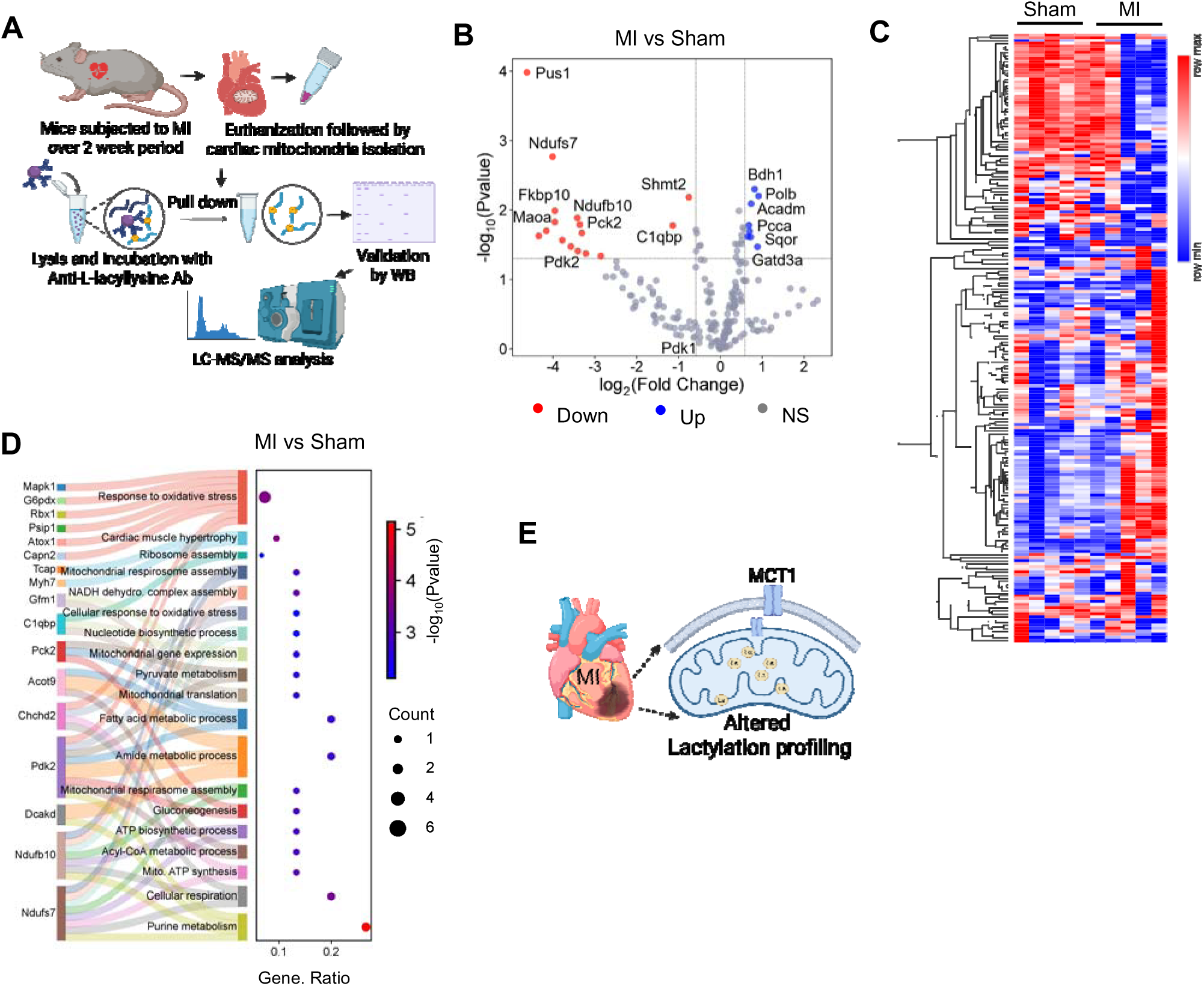
MI remodels the mitochondrial lactylome. (A) Schematic overview of the experimental workflow for mitochondrial lactylome profiling in mouse hearts using LC-MS/MS. (B) Volcano plot showing significantly altered mitochondrial protein lactylation in MI versus sham hearts. (C) Heatmap visualization of differentially lactylated mitochondrial proteins in MI versus sham hearts. Hierarchical clustering was used to group proteins with similar lactylation patterns. (D) Sankey dot plot illustrating Gene Ontology (GO) enrichment analysis of biological processes associated with mitochondrial proteins exhibiting significantly altered lactylation (P < 0.05). Dot size represents the number of hyper or hypo-lactylated proteins within each GO term. (E) Schematic summary depicting mitochondrial protein lactylation remodeling following myocardial infarction in mice. Lactylome data are derived from n = 5 mice per group.

Gene ontology (GO) enrichment analysis of differentially lactylated mitochondrial proteins revealed significant enrichment in biological processes related to oxidative stress response, protein folding, RNA splicing, cardiac muscle growth, translation, oxidative phosphorylation, and cellular respiration, as well as pathways associated with cardiac hypertrophy and muscle adaptation (Fig. 1D). These results indicate that MI-induced changes in mitochondrial lactylation preferentially affect proteins involved in metabolic regulation, mitochondrial function, and cardiac remodeling. To determine whether lactylation occurs within mitochondrial compartments rather than being restricted to the outer mitochondrial membrane, isolated mitochondria from AC16 (human cardiomyocytes derived) cells were subjected to limited proteolysis using proteinase K under intact conditions. While proteinase K efficiently degraded proteins exposed on the outer mitochondrial membrane, global lactylation signals were largely preserved, displaying patterns comparable to untreated mitochondria (Fig. S1D, E). These findings indicate that lactylation is present within protease-protected mitochondrial compartments, including the intermembrane space and matrix. Collectively, these data demonstrate that myocardial infarction induces extensive remodeling of the mitochondrial lactylome, affecting a diverse set of proteins involved in mitochondrial metabolism, redox regulation, and cardiac structural adaptation (Fig. 1E).

### MCT1 inhibition reshapes the mitochondrial lactylome following MI

To determine whether inhibition of lactate transport alters mitochondrial protein lactylation following myocardial infarction, AZD3965 was administered to MI mice and mitochondrial lactylome profiling was performed. Heatmap visualization revealed a pronounced remodeling of mitochondrial protein lactylation in the MI + AZD3965 group compared with sham and MI alone (Fig. 2A). Volcano plot analysis further delineated proteins exhibiting significant hyper-and hypo-lactylation across experimental groups (Fig. 2B, C). In the MI + AZD3965 versus sham comparison, lactylation was significantly increased in 33 mitochondrial proteins, including ACADM, BDH1, IDH3A, POLB, ECHS1, MGST3, HADHA, PCCA, SLC25A5, CPT1B, ACADS, NDUFA10, GATD3A, HSDL2, COQ8A, SLC25A4, CKMT2, OGDH, TUFM, SLC25A12, UQCRC1/2, HADHB, ATP5PB, VDAC2, DECR1, HSD17B10, ACOT13, and CS (Fig. 2B). In contrast, significant reductions in lactylation were observed in 20 proteins, including FASN, SHMT2, ALDH18A1, UQCRB, MRPL12, HINT1, PUS1, HSPE1, COX7A2, GPD2, NDUFB10, PCK2, APEX1, PRDX6, ALDH2, and SLC25A4 (Fig. 2B). Comparison of MI + AZD3965 versus MI alone revealed increased lactylation of 26 proteins, including NDUFS7, IDH3A, CS, ECHS1, SUCLG1, HSDL2, HSD17B10, ACADS, NDUFA10, ACADM, ACOT2, PDK2, PDHX, DLD, HADHA, BDH1, COQ8A, NNT, and GSTK1, while 16 proteins—including UQCRB, CHCHD3, NDUFB8, MRPL12, SOD2, PRDX2, COX7A2, FASN, PRDX4, GPD2, CKMT1, SUCLG2, and COX5A, displayed reduced lactylation (Fig. 2C).

**Figure 2.**
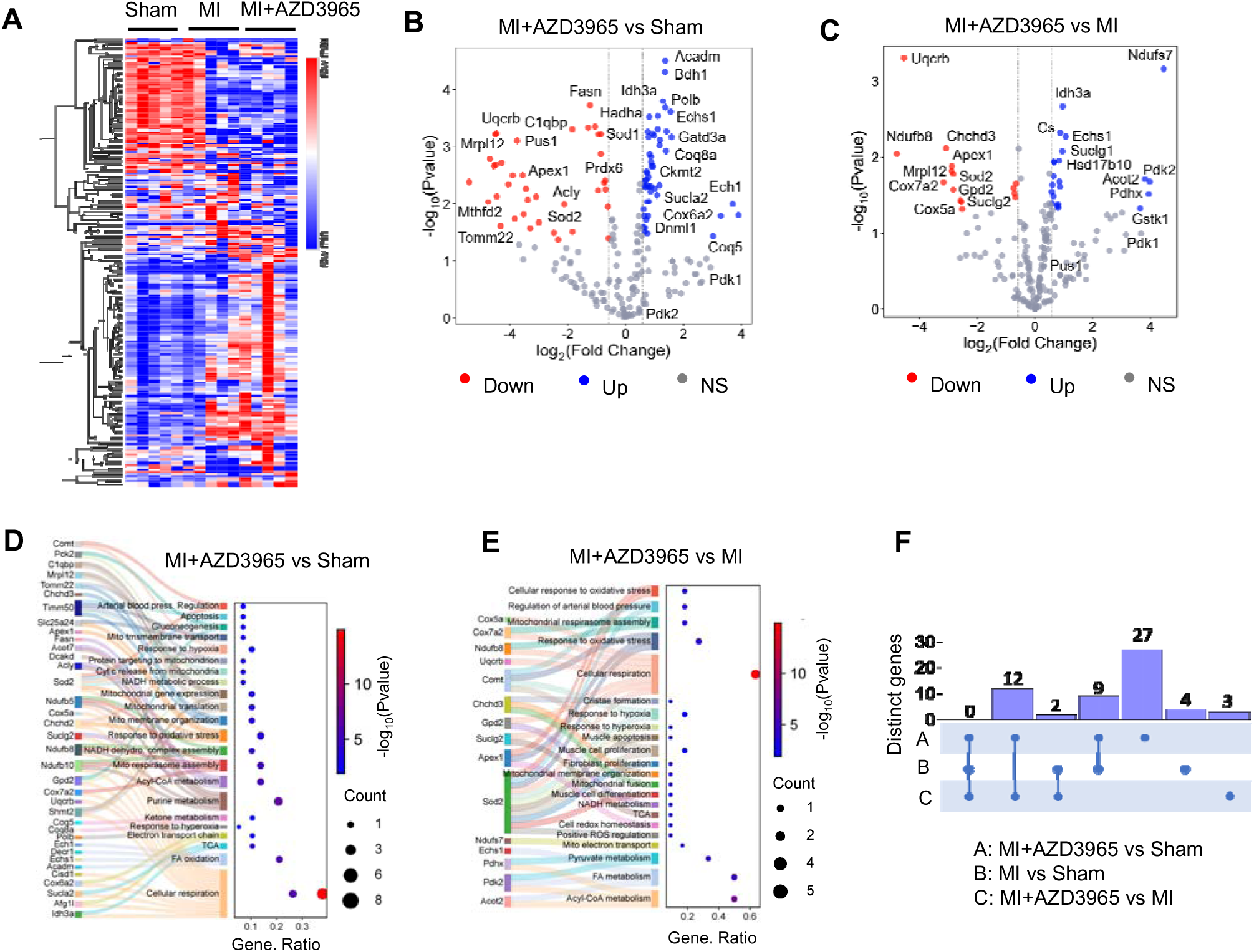
MCT1 inhibition reshapes the mitochondrial lactylome following MI. (A) Heatmap illustrating significantly altered mitochondrial protein lactylation across sham, MI, and MI + AZD3965 groups in mouse hearts two weeks after MI. Hierarchical clustering was used to group proteins with similar lactylation patterns. n = 5 mice per group. (B, C) Volcano plots showing significantly altered mitochondrial protein lactylation in MI + AZD3965 versus sham (B) and MI + AZD3965 versus MI (C). (D, E) Sankey dot plots depicting GO enrichment analysis of biological processes associated with mitochondrial proteins exhibiting significantly altered lactylation in MI + AZD3965 versus sham (D) and MI + AZD3965 versus MI (E). (F) UpSet plot showing the overlap among sets of differentially lactylated mitochondrial proteins identified in pairwise comparisons (MI + AZD3965 vs sham, MI vs sham, and MI + AZD3965 vs MI). Filled dots indicate inclusion of a comparison in the intersection, and vertical bars represent the number of proteins in each intersecting set.

GO enrichment analysis was visualized using Sankey dot plots [29] to illustrate biological processes associated with differentially lactylated proteins (Fig. 2D, E; Fig. S1F). In the MI + AZD3965 versus sham comparison, proteins exhibiting increased lactylation were enriched in processes related to fatty acid β-oxidation, lipid catabolism, tricarboxylic acid cycle, lipid oxidation, and mitochondrial translation, whereas proteins with reduced lactylation were enriched in processes including cellular respiration, oxidative phosphorylation, succinyl-CoA metabolism, oxidative stress response, and mitochondrial membrane organization (Fig. 2D). Relative to MI alone, MI + AZD3965 samples showed enrichment of hyper-lactylated proteins associated with acetyl-CoA metabolism, fatty acid metabolism, thioester metabolism, and oxidant detoxification, while hypo-lactylated proteins were enriched in pathways related to cellular respiration, ATP synthesis, mitochondrial membrane organization, mitofusion, redox homeostasis, and the TCA cycle (Fig. 2E).

Notably, lactylation of PDK2, GSTK1, and PDHX was reduced in MI versus sham but increased in MI + AZD3965 versus MI, indicating dynamic regulation of these proteins by lactate transport inhibition. Intersection analysis using UpSet plots revealed both unique and shared sets of differentially lactylated proteins across MI + AZD3965 versus sham, MI versus sham, and MI + AZD3965 versus MI comparisons, with the largest subset (27 proteins) unique to the MI + AZD3965 versus sham group (Fig. 2F; Table S3). Collectively, these data demonstrate that MCT1 inhibition markedly reshapes the mitochondrial lactylome following myocardial infarction, affecting proteins involved in mitochondrial metabolism, redox regulation, and energy production.

### MCT1 blockade attenuates cardiac inflammation and fibrosis following MI

To evaluate the impact of MCT1 inhibition on cardiac remodeling following myocardial infarction, cardiac function was assessed by echocardiography at baseline, and at one and two weeks post-MI in sham, MI, and MI + AZD3965 groups. Left ventricular systolic function was significantly impaired in both MI and MI + AZD3965 mice compared with sham controls, as reflected by reduced ejection fraction and fractional shortening at one and two weeks post-MI (Fig. 3A-C). No significant differences in these parameters were observed between MI and MI + AZD3965 groups during the study period. Plasma levels of B-type natriuretic peptide (BNP), a marker of ventricular wall stress [32], showed an upward trend in MI mice compared with sham, although this increase did not reach statistical significance (Fig. 3D). Notably, BNP levels were significantly lower in MI + AZD3965 mice compared with MI mice (Fig. 3D). Consistent with these findings, the heart weight-to-tibia length ratio was significantly increased in MI mice relative to sham, whereas MI + AZD3965 mice exhibited values comparable to sham controls (Fig. S1C).

**Figure 3.**
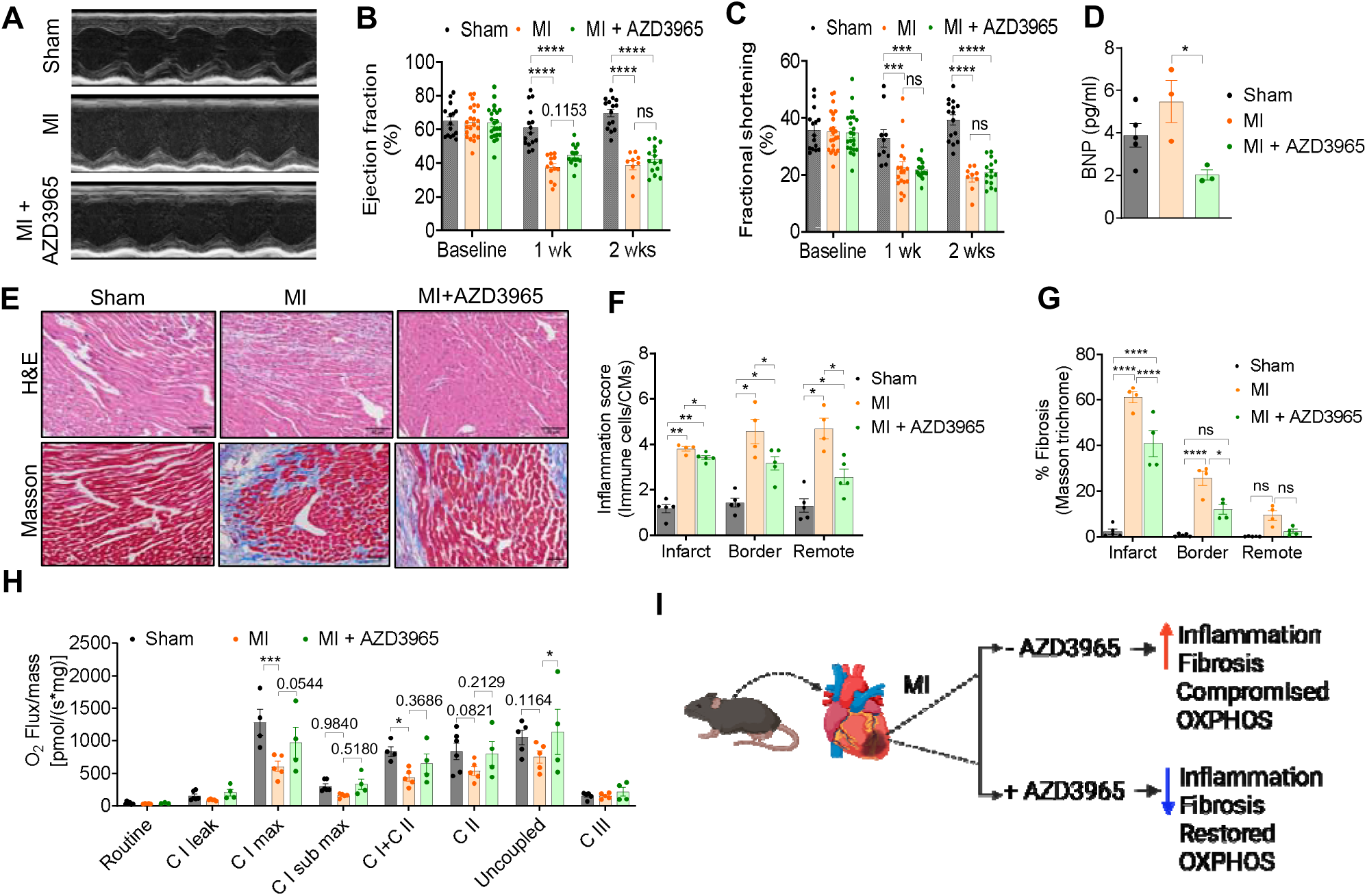
MCT1 blockade attenuates cardiac inflammation and fibrosis following MI. (A) Representative M-mode echocardiographic images from sham, MI, and MI + AZD3965 mice two weeks post-MI. (B, C) Quantification of echocardiographic parameters, including ejection fraction (B) and fractional shortening (C), in sham, MI, and MI + AZD3965 mice two weeks after sham or MI surgery (n = 12–15 mice per group). (D) Plasma BNP levels measured by ELISA (n = 4–5 mice per group). Statistical significance was determined by one-way ANOVA with Tukey’s multiple-comparisons test. (E) Representative images of H&E and Masson’s trichrome staining from sham, MI, and MI + AZD3965 hearts collected 14 days after sham or MI surgery. Scale bar, 50 µm (n = 4–5 mice per group). (F, G) Quantification of cardiac inflammation score (F) and fibrotic area (G) derived from histological analyses shown in E (n = 4–5 mice per group). Statistical significance was determined by one-way ANOVA with Tukey’s multiple-comparisons test. (H) Mitochondrial respiration was measured using the Oroboros O2k system in permeabilized heart tissue. Routine respiration and substrate– and inhibitor-specific respiratory states were assessed, including CI-linked respiration (pyruvate + malate), CI maximal respiration (ADP + Glutamate), CI sub-maximal respiration (ADP),CI + CII respiration (glutamate + succinate), CII-linked respiration (succinate), uncoupled respiration (FCCP), and residual respiration following CIII inhibition (antimycin A) (n = 4–5 mice per group). Statistical significance was determined by two-way ANOVA with Sidak’s multiple-comparisons test. (I) Schematic summary illustrating the effects of MCT1 inhibition on cardiac remodeling following myocardial infarction. Data are presented as mean ± SEM. P < 0.05, P < 0.01, P < 0.001, P < 0.0001.

Histological assessment using H&E staining revealed prominent inflammatory cell infiltration and myocardial tissue disruption in MI hearts compared with sham hearts (Fig. 3E). Quantitative analysis demonstrated a significant reduction in inflammation scores in MI + AZD3965 hearts relative to MI, approaching levels observed in sham mice (Fig. 3F). Because fibrosis is a hallmark of adverse cardiac remodeling following MI, fibrotic burden was assessed by Masson’s trichrome and Sirius red staining across infarct, border, and remote zones. Both staining methods revealed marked increases in fibrosis within the infarct and border zones of MI hearts compared with sham controls (Fig. 3E, G; Fig. S1G, H). While MI + AZD3965 hearts exhibited increased fibrosis relative to sham, fibrotic area in the infarct and border zones was significantly reduced compared with MI hearts (Fig. 3E, G; Fig. S1G, H).

Mitochondrial respiratory function was assessed to determine whether MCT1 inhibition influences mitochondrial performance following MI. Mitochondrial respiration was impaired in MI hearts compared with sham, whereas MI + AZD3965 hearts exhibited a modest but reproducible improvement in respiratory capacity (Fig. 3H). MI hearts showed a significant reduction in complex II (succinate) linked respiration, which was restored in MI + AZD3965 hearts (Fig. 3H), compensating for the compromised respiration[33]. Collectively, these findings indicate that MCT1 inhibition via AZD3965 attenuates cardiac inflammation and fibrosis following MI, while having a limited impact on global systolic function during the early post-infarction period (Fig. 3I).

### MCT1 inhibition remodels mitochondrial lactylation and attenuates profibrotic and pro-inflammatory signaling in hypoxic AC16 cells

To examine the role of MCT1 in regulating mitochondrial lactylation under hypoxic stress, AC16 cells were subjected to hypoxia following either genetic (siRNA-mediated knockdown) or pharmacological (AZD3965) inhibition of MCT1 (Fig. 4A). Efficient knockdown of MCT1 was confirmed by immunoblotting (Fig. S2A, B). Under hypoxic conditions, suppression of MCT1 by either approach resulted in a modest but non-significant increase in global mitochondrial protein lactylation, as assessed by pan-lactyllysine immunoblotting (Fig. S2C, D). To determine whether specific mitochondrial metabolic regulators are selectively lactylated, we performed immunoprecipitation of pyruvate dehydrogenase kinases PDK1 and PDK2. Hypoxia markedly increased lysine lactylation of both PDK1 and PDK2 compared with normoxia (Fig. 4B–D). In contrast, neither MCT1 knockdown nor AZD3965 treatment further altered PDK1 or PDK2 lactylation under hypoxic conditions, indicating that hypoxia alone is sufficient to induce maximal lactylation of these kinases.

**Figure 4.**
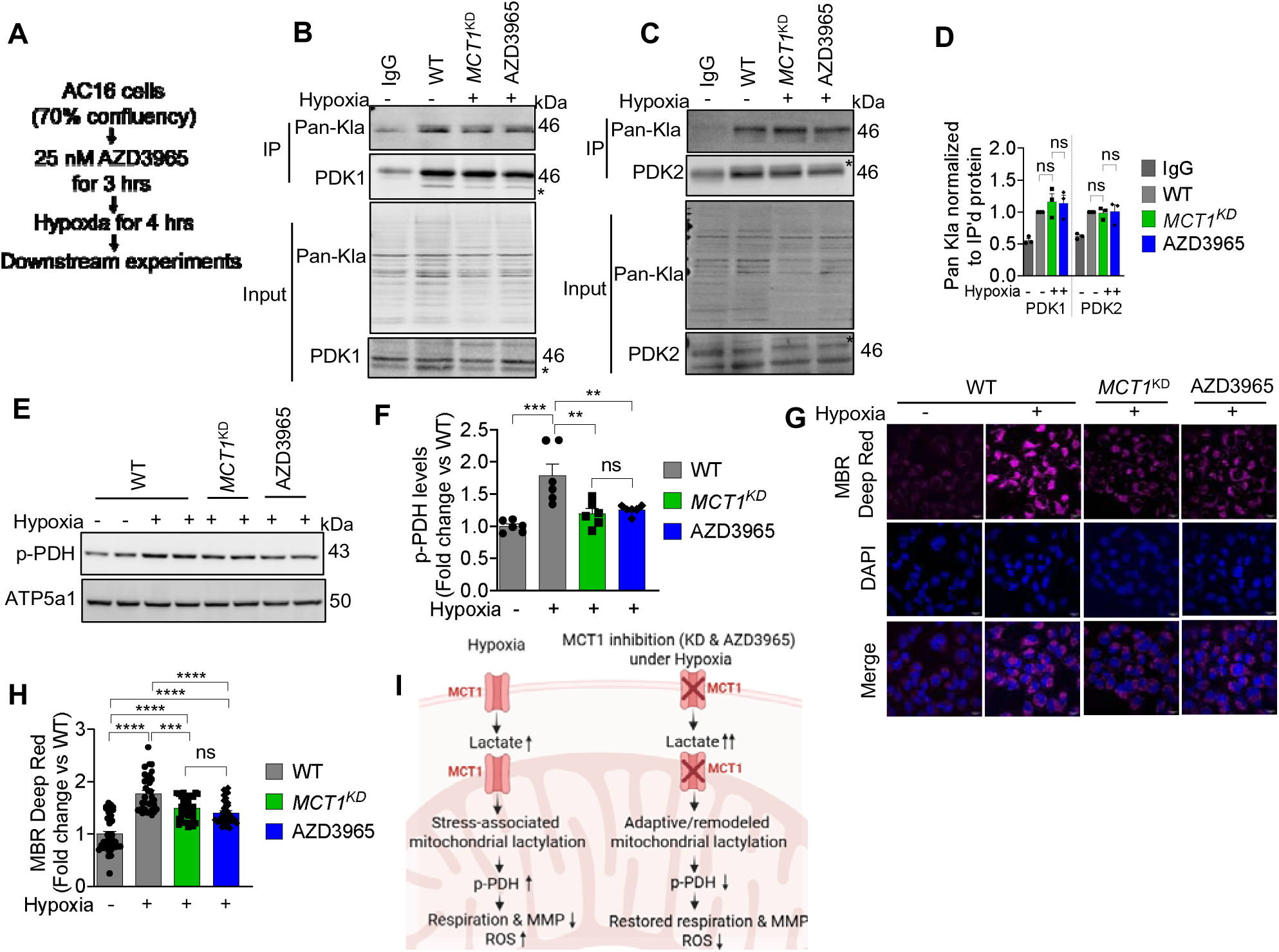
MCT1 inhibition reprograms mitochondrial lactylation and mitochondrial function in hypoxic AC16 cells. (A) Schematic overview of the experimental workflow and timeline for genetic (siRNA-mediated knockdown) and pharmacological (AZD3965) inhibition of MCT1 in AC16 cardiomyocytes-derived cells under hypoxic conditions. (B, C) Immunoblot analysis of lysine lactylation on PDK1 (B) and PDK2 (C) following immunoprecipitation from AC16 cells under normoxic or hypoxic conditions with or without MCT1 knockdown or AZD3965 treatment. (D) Quantification of PDK1 and PDK2 lactylation normalized to immunoprecipitated protein levels. (E) Representative immunoblot showing p-PDH levels in AC16 cells subjected to hypoxia with genetic or pharmacological inhibition of MCT1. (F) Quantification of p-PDH protein levels. (G) Representative confocal images showing mitochondrial ROS generation detected using MitoBright ROS Deep Red (MBR Deep Red) in hypoxic AC16 cells with MCT1 knockdown or AZD3965 treatment. (H) Quantification of mitochondrial ROS fluorescence intensity. Scale bar, 20 µm; magnification, 60x. (I) Schematic summary illustrating MCT1-dependent modulation of mitochondrial lactylation, PDH regulation, and mitochondrial function under hypoxic conditions in AC16 cells. Data are presented as mean ± SEM from n = 3–6 independent experiments. Statistical significance was determined by one-way ANOVA with Tukey’s multiple-comparisons test. P < 0.05, P < 0.01, P < 0.001, P < 0.0001.

Given the central role of PDKs in regulating pyruvate dehydrogenase (PDH) activity, we next assessed PDH phosphorylation status. Hypoxia significantly increased phosphorylation of PDH at inhibitory sites, consistent with reduced PDH activity. Notably, both MCT1 knockdown and AZD3965 treatment significantly reduced PDH phosphorylation under hypoxia (Fig. 4E, F), indicating modulation of PDH activity independent of further changes in PDK lactylation. To evaluate the functional consequences of MCT1-dependent metabolic remodeling, mitochondrial respiration, ROS production, and mitochondrial membrane potential were assessed. Hypoxia significantly impaired mitochondrial respiration, whereas both genetic and pharmacological inhibition of MCT1 restored respiratory capacity (Fig. S3A, B). Hypoxia also induced a robust increase in mitochondrial ROS production, which was significantly attenuated by MCT1 knockdown or AZD3965 treatment (Fig. 4G, H). To further characterize mitochondrial redox remodeling, we examined lactylation changes in mitochondrial proteins associated with oxidative stress regulation and redox homeostasis.

Heatmap analysis revealed distinct patterns of lactylation in redox-related mitochondrial proteins under hypoxic conditions, which were altered following genetic or pharmacological inhibition of MCT1 (Fig. S3C). In parallel, mitochondrial membrane potential was restored in MCT1-inhibited hypoxic cells compared with hypoxia alone (Fig. S3D, E). Collectively, these results indicate that hypoxia induces near maximal lactylation of mitochondrial PDKs, while MCT1 inhibition reshapes downstream mitochondrial metabolism and redox balance without further altering PDK lactylation, suggesting the involvement of additional lactylation-dependent mitochondrial targets (Fig.□4I). Thus, the functional impact of mitochondrial lactylation depends on protein targets and metabolic context rather than absolute lactylation levels.

To determine whether MCT1 inhibition also influences profibrotic and pro-inflammatory signaling under hypoxic conditions, transcript levels of key markers were assessed by quantitative real-time PCR. Hypoxia significantly increased expression of fibrosis-associated genes ACTA2 and FN1, as well as inflammatory markers IL6 and TNFα, compared with normoxic controls (Fig. S4A, B). Notably, both genetic and pharmacological inhibition of MCT1 significantly attenuated hypoxia-induced expression of these genes, restoring transcript levels toward baseline. These findings are consistent with *in vivo* observations and indicate that MCT1 inhibition suppresses hypoxia-induced profibrotic and pro-inflammatory signaling in cardiomyocytes.

## Discussion

Lactate has evolved from being viewed solely as a glycolytic byproduct to a key metabolic and signaling intermediate, as exemplified by the Warburg effect and emerging roles of protein lactylation [5, 7, 34, 35]. Recent studies have implicated lactylation in cardiovascular pathophysiology, influencing inflammation, fibrosis, and tissue remodeling, and have highlighted lactate transporters and lactylation as potential therapeutic targets in cardiovascular disease [36, 37]. Notably, Wang et al. demonstrated that lactylation of Serpina3k enhances its stability and secretion from cardiac fibroblasts, conferring cardioprotection following ischemic injury [38]. Building on these findings, our study investigated how MI reshapes the mitochondrial lactylome and whether modulation of lactate transport via MCT1 influences this process. Using quantitative proteomics, we demonstrate that MI induces extensive remodeling of mitochondrial protein lactylation, with both hypo-and hyper-lactylation affecting proteins involved in energy metabolism, redox regulation, and mitochondrial organization. Several lactylated proteins identified here, including VDAC isoforms, HADHA, BDH1, and PCK2, have previously been implicated in cardiac or hypoxia-associated pathologies [39–43], whereas others represent novel candidates linking mitochondrial lactylation to MI. Importantly, many of these proteins have not previously been examined in the context of lactylation, underscoring the exploratory and hypothesis generating nature of our dataset.

Pharmacological inhibition of MCT1 with AZD3965 further reshaped the mitochondrial lactylome in both in vivo MI and hypoxic cardiomyocyte models. Given the bidirectional nature of MCT1-mediated lactate transport, MCT1 inhibition likely promotes intracellular lactate accumulation and redistribution rather than simple deprivation of mitochondrial lactate. Consistent with this model, both genetic and pharmacological MCT1 inhibition produced comparable alterations in mitochondrial lactylation patterns, supporting a central role for MCT1 in controlling mitochondrial lactate availability and downstream post-translational modification. Mitochondria can produce lactate themselves, even when energized, rather than preferentially oxidizing it [8]. This also could be one of the reasons for increased protein lactylation upon MCT1 inhibition. Despite extensive lactylome remodeling, MCT1 inhibition did not significantly improve global cardiac function within the studied timeframe. This likely reflects the multifactorial nature of post-MI remodeling, where modulation of a single metabolic axis may be insufficient to elicit rapid organ-level functional recovery. Nevertheless, MCT1 inhibition partially attenuated cardiac inflammation and fibrosis, and reduced expression of fibrotic and pro-inflammatory genes in hypoxic cardiomyocytes, suggesting that mitochondrial lactylation may influence reparative and inflammatory signaling rather than acute contractile performance.

At the mitochondrial level, MCT1 inhibition paradoxically enhanced mitochondrial protein lactylation while improving respiratory capacity, membrane potential, and redox balance. These findings indicate that the functional consequences of mitochondrial lactylation are highly context and target-dependent. While hypoxia-induced lactylation is associated with mitochondrial dysfunction, lactylation occurring in the setting of altered lactate transport may instead support adaptive metabolic remodeling. In this context, improved mitochondrial respiration and reduced ROS production may reflect lactylation of non-PDK targets, including components of the PDH complex, PDH phosphatases, or electron transport chain subunits, rather than changes in PDK lactylation per se. Consistent with this interpretation, MCT1 inhibition reduced PDH phosphorylation under hypoxia, suggesting restoration of pyruvate oxidation despite unchanged PDK lactylation. This dissociation highlights that lactylation does not act as a uniform inhibitory signal, but rather integrates with lactate transport, redox state, and substrate utilization to shape mitochondrial function. Similar metabolic adaptations have been reported in skeletal muscle, where MCT1 deletion enhances mitochondrial biogenesis and oxidative capacity through NAD⁺-SIRT1-PGC-1α signaling, supporting a conserved role for MCT1 in regulating mitochondrial metabolism across tissues [44].

Collectively, our findings suggest that mitochondrial lactylation serves as an adaptive regulatory signal rather than merely a pathological byproduct of elevated lactate. However, excessive or dysregulated lactylation, particularly in processes such as fibrosis and endothelial-to-mesenchymal transition, may contribute to adverse cardiac remodeling, highlighting the dual nature of this modification. Thus, therapeutic strategies targeting lactate transport or lactylation must consider disease stage, cell type, and metabolic context to balance beneficial and detrimental effects.

In conclusion, our study reveals a previously underappreciated link between mitochondrial protein lactylation, lactate transport, and cardiac remodeling following MI. By reshaping the mitochondrial lactylome and improving mitochondrial metabolic efficiency, MCT1 inhibition partially mitigates post-MI pathology. These findings position AZD3965 as a valuable tool compound for probing lactate-dependent mitochondrial signaling and lay the foundation for future studies aimed at therapeutically modulating metabolic plasticity in ischemic heart disease.

## Author Contributions

**AK**: Investigation, validation, formal data analysis, writing-original draft, methodology.

**SK, KS, NJ, SG, PK, PH, JL, and CF:** Formal data analysis, resources, and methodology.

**DT, and PJ**: Conceptualization, methodology, resources, visualization, supervision, writing – review and editing, funding acquisition.

## Acknowledgments

We thank both past and present members of the Tomar and Jadiya labs for their valuable support, scientific input, methodological assistance, and resources. We acknowledge Core for Cellular Respirometry, Tumor Tissue and Pathology Shares Resource and Cellular Imaging Shared Resource at Wake Forest University School of Medicine for assistance with mitochondrial bioenergetics, histology and microscopy. The language editing was completed using Microsoft Co-pilot. Schematics were created with BioRender.com.

## Data availability

The mass spectrometry proteomics data have been deposited to the ProteomeXchange Consortium via the PRIDE partner repository with the dataset identifier PXD068087. Raw proteomics data associated with this manuscript can be accessed by following credentials: Reviewer access details- Project accession: “PXD068087”, Token: “cRIUWyxJVEsP”. The rest of the raw data deposited to figshare and can be accessed using the DOI: 10.6084/m9.figshare.32035257.

