## Supplementary material for "Lactylation landscape of mitochondrial proteins in myocardial infarction": Fig. S

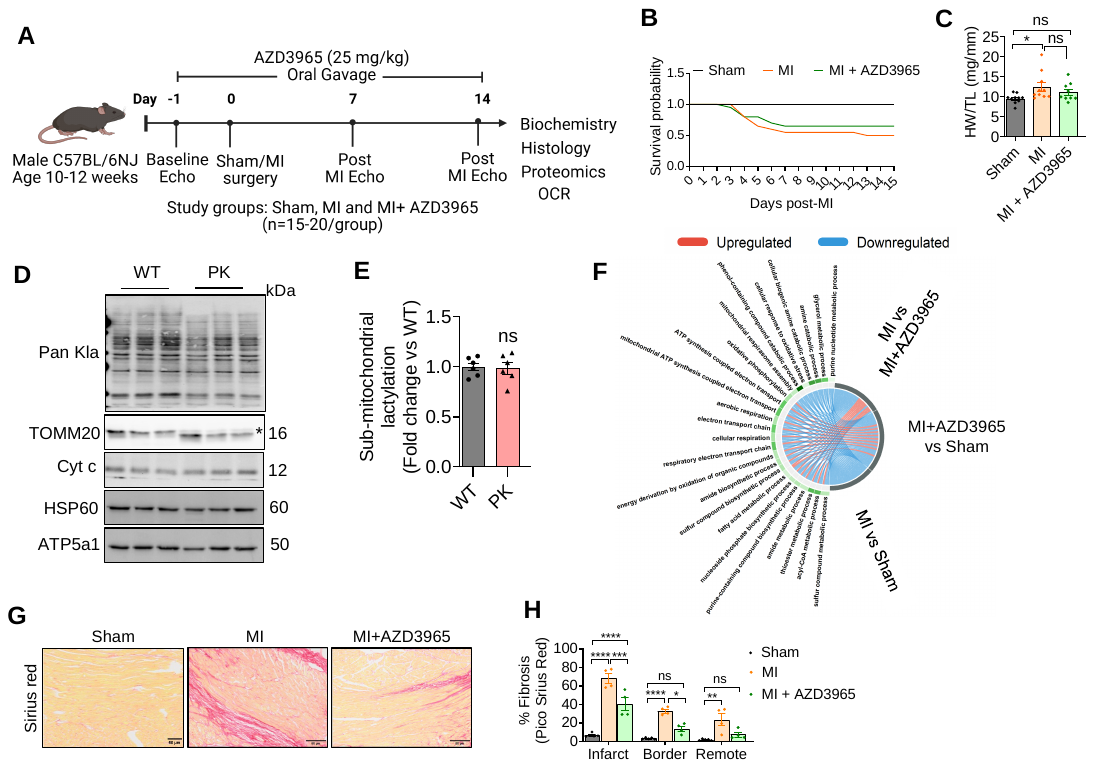
**Supplementary Data**

**Supplementary Figure 1: MI remodels the mitochondrial lactylome profiling**

A. Systemic workflow of the experimental timeline for MI and AZD3965 administration for the study. B. Survival curve plot after sham and MI surgeries, analyzed over two weeks. C. Ratios of heart weight to body weight (n=5–10). D. Mitochondrial fractions from WT AC16 cells were subjected to Proteinase K treatment (3 µg/ml) to remove OMM-exposed proteins and probed with mitochondria compartment-specific indicated antibodies. Western blots are representative of six independent replicates. E. Quantification of submitochondrial lactylation (n=6); ns: non-significant. F. The chord diagram represents the relationship between the GO term and lactylated proteins across the sham, MI, and MI + AZD3965 groups. G. Representative images of Pico Sirius red staining from sham, MI, and MI+AZD3965 mice post-sham or MI surgery. Scale bar, 50 µm, n=4-5 mice in each group. H. Quantification of cardiac fibrosis was quantified from D. n=4-5 mice in each group.

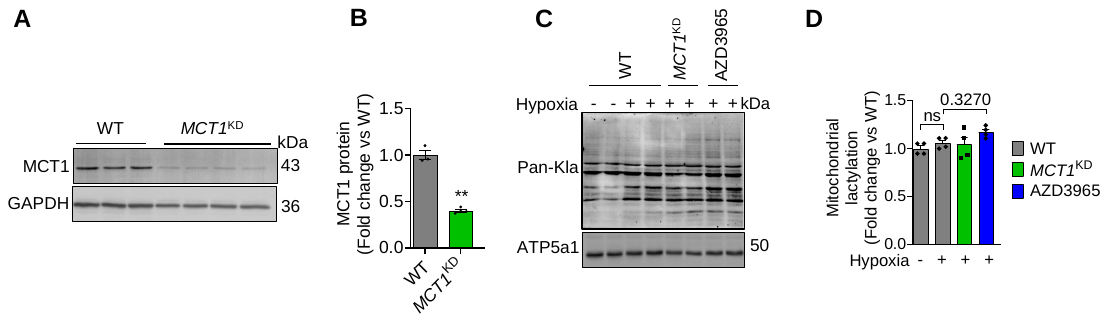

**Supplementary Figure 2: MCT1 inhibition reprograms the mitochondrial lactylome in hypoxic AC16 cells**

A. Western blot image showing characterization of siRNA-mediated *MCT1* knockdown in AC16 cells. B. Quantification of MCT1 protein normalized to GAPDH protein levels. C. Western blot image showing lactylation of mitochondrial proteins in hypoxic *MCT1*^KD^ and AZD3965-treated cells. D. Quantification of mitochondrial protein lactylation. Data are presented as mean ± SEM from n = 3-4 independent experiments. Statistical significance was determined by Unpaired t-test with Welch’s correction and one-way ANOVA with Tukey’s multiple comparisons tests. **p < 0.01.

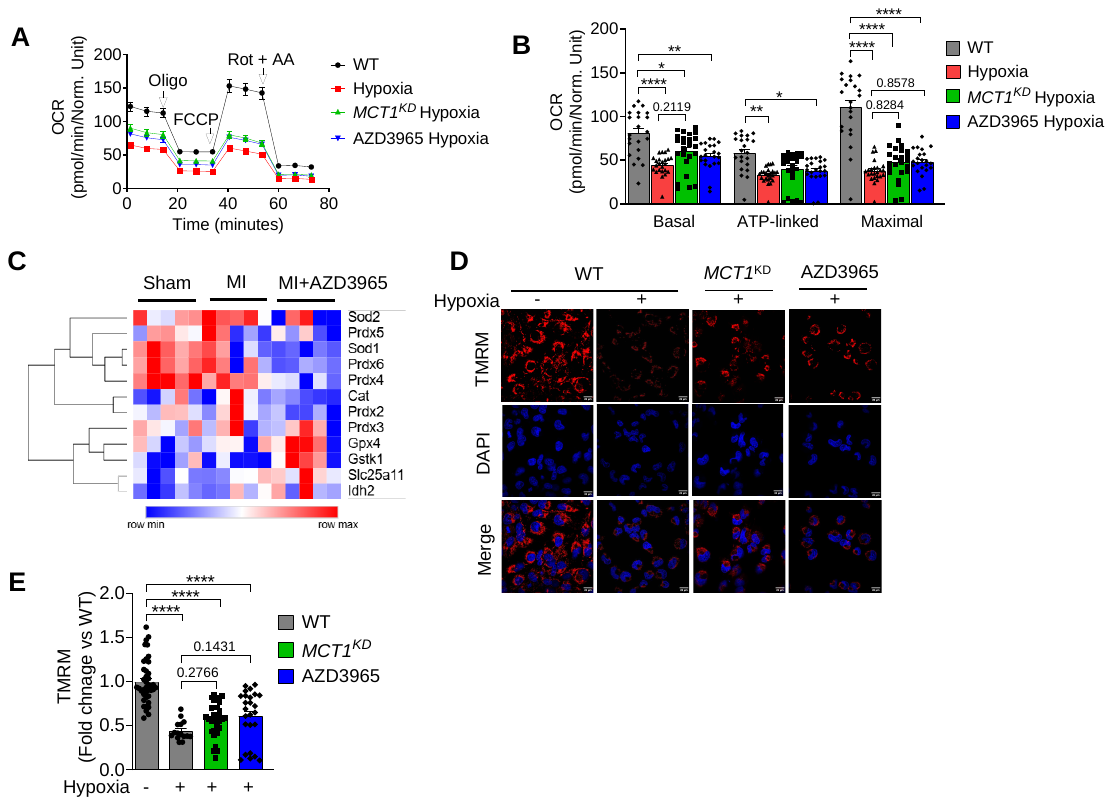

**Supplementary Figure 3: MCT1 inhibition improves mitochondrial function under hypoxia in AC16 cells**

A. Representative traces of oxygen consumption rate (OCR) in hypoxic *MCT1*^KD^ and AZD3965-treated cells at baseline and following: oligomycin (oligo; Complex V inhibitor; to uncover ATP-linked respiration), FCCP (protonophore to induce maximum respiration), and rotenone + antimycin A (Rot/AA; complex I and III inhibitor for complete ETC inhibition). B. Quantification of OCR. C. A heat map shows the lactylation levels of ROS-related proteins.

D. Representative images of mitochondrial membrane potential using TMRM dye in hypoxic *MCT1*^KD^ and AZD3965-treated cells. E. Quantification of mitochondrial membrane potential. Scale bar 20 µm; Magnification 60x. Data are presented as mean ± SEM from n = 3 independent experiments. Statistical significance was determined by one-way ANOVA with Tukey’s multiple comparisons test. *p < 0.05, **p < 0.01, ****p < 0.0001.

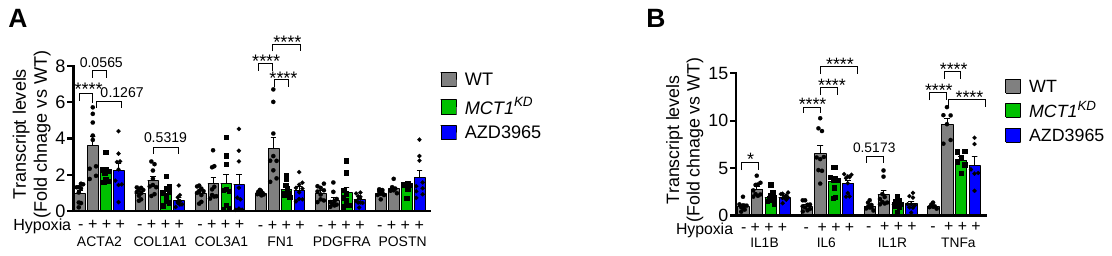

**Supplementary Figure 4: MCT1 inhibition remodels the cardiac fibrosis in hypoxic AC16 cells**

A-B. Analysis of transcript levels of candidate genes of fibrosis (A) and inflammation (B) signalling by real-time PCR in hypoxic AC16 cells upon MCT1 inhibition. Data are presented as mean ± SEM from n = 3 independent experiments. Statistical significance was determined by Unpaired t-test with Welch’s correction and one-way ANOVA with Tukey’s multiple comparisons tests. ****p < 0.0001.

**Supplementary Table 1:** The list of primer sequences used in the study.

| **Gene**  **(Human)** | **Forward primer** | **Reverse primer** |
| --- | --- | --- |
| *GAPDH* | GGAGCGAGATCCCTCCAAAAT | GGCTGTTGTCATACTTCTCATGG |
| *ACTIN* | CACCATTGGCAATGAGCGGTTC | AGGTCTTTGCGGATGTCCACGT |
| *ACTA2* | CGACCGAATGCAGAAGGA | ACAGAGTATTTGCGCTCCGAA |
| *COL1A1* | TCTGCGACAACGGCAAGGTG | GACGCCGGTGGTTTCTTGGT |
| *COL3A1* | TTCTCGCTCTGCTTCATCCC | TCCGCATAGGACTGACCAAG |
| *FN1* | CGGTGGCTGTCAGTCAAAG | AAACCTCGGCTTCCTCCATAA |
| *PDGFRA* | GGGCACGCTCTTTACTCCAT | TTAGGCTCAGCCCTGTGAGA |
| *POSTN* | CCATGTTTATGGCACTCTGG | ACGTTGCTCTCCAAACCTCT |
| *IL1B* | ATGCACCTGTACGATCACTG | ACAAAGGACATGGAGAACACC |
| *IL6* | CCACTCACCTCTTCAGAACG | CATCTTTGGAAGGTTCAGGTTG |
| *IL1R* | GATGAAGATGACCCAGTGCTAG | TGGCAAAACAGGTAAATGGATG |
| *TNFa* | ACTTTGGAGTGATCGGCC | GCTTGAGGGTTTGCTACAAC |

**Supplementary Table 2: Predicted Lysine lactylation sites in differentially expressed proteins in MI vs Sham:** The table lists the predicted lysine lactylation sites in proteins that were significantly differentially expressed between MI vs Sham group. Lactylation site prediction was performed using the DeepKla software.

| **Sr no.** | **Name of the protein** | **Lactylation sites (>50 %)** |
| --- | --- | --- |
| 1 | CKMT1 | K292, K296, K344, K353 |
| 2 | PPA2 | K254, K256, K276, K280 |
| 3 | IDH3G | K206 |
| 4 | GSTK1 | K49, K68, K85, K94, K165, K167, K177 |
| 5 | PUS1 | K76, K79, K81, K180, K189, K191, K292, K302 |
| 6 | AHCYL2 | K136, K180, K191, K194, K350, K353, K367, K472 |
| 7 | AHCYL1 | K36, K39, K40, K53, K108, K111, K267, K270, K284, K286, K302, K389, |
| 8 | PDHX | K213, K218, K223, K227, K394, K398, K399 |

**Supplementary Table 3: Intersections of differentially expressed proteins across experimental comparisons:** The table summarizes the protein details by the intersection as shown in the UpSet plot they belong to (e.g., shared across all comparisons, unique to one, or shared between two)

| **Intersection set** | **Name of proteins shared** | **No. of counts shared** |
| --- | --- | --- |
| All three group comparisons | NA | 0 |
| MI+AZD3965 vs Sham and MI+AZD3965 vs MI | APEX1, CHCHD3, COMT, COX5A, COX7A2, ECHS1, GPD2, MRPL12, NDUFB8, SOD2, SUCLG2, UQCRB | 12 |
| MI vs Sham and MI+AZD3965 vs MI | NDUFS7, PDK2 | 2 |
| MI+AZD3965 vs Sham and MI vs Sham | C1QBP, CHCHD2, DCAKD, MAOA, MTHFD2, NDUFB10, PCK2, PUS1, TOMM22 | 9 |
| Unique to MI+AZD3965 vs Sham | ACADM, ACLY, ACOT7, AFG1L, ALDH18A1, BDH1, CISD1, COQ5, COQ8A, COX6A2, DBI, DECR1, ECH1, FASN, GATD3A, HSDL2, IDH3A, MCCC1, MGST3, NDUFB5, PCCA, POLB, SHMT2, SLC25A24, SUCLA2, TIMM50, VWA8 | 27 |
| Unique to MI vs Sham | ACOT9, CYB5B, FKBP10, GFM1 | 4 |
| Unique to MI+AZD3965 vs MI | ACOT2, GSTK1, PDHX | 3 |
